# Climate change, age acceleration, and the erosion of fitness in polar bears

**DOI:** 10.1101/2024.01.05.574416

**Authors:** Levi Newediuk, Evan S Richardson, Brooke A. Biddlecombe, Haziqa Kassim, Leah Kathan, Nicholas Lunn, L Ruth Rivkin, Ola E Salama, Chloé Schmidt, Meaghan J Jones, Colin J Garroway

## Abstract

Climate change is increasingly disrupting evolved life history strategies and reducing population viability in wild species. Using estimates of epigenetic age acceleration, a cellular biomarker of lifetime stress and the expression of age-related phenotypes, we found that polar bears aged approximately one year faster for each degree of warming since the 1960s. Age acceleration was also associated with reproducing early in life, linking this cellular process to well-established life history theory. However, we found evidence for the erosion of fitness as epigenetic aging accelerated and temperatures increased. Finally, using a large pedigree, we found adaptive potential in our study population was approximately zero. Global temperatures will soon reach the levels of warming currently experienced by Arctic species, which could impose widespread physiological costs and limit adaptive capacities worldwide.

## Introduction

Climate change is causing extreme environmental fluctuations and persistent warming, exposing species to conditions increasingly distant from those they evolved to tolerate(*1*). Species have typically responded to changes in climate through range shifts(*2*), changes in seasonally timed behaviours(*3*), and population declines(*4*). However, the accumulation of our past emissions and current emission targets commits the planet to ongoing warming for the foreseeable future(*5*). This means the persistence of many species will ultimately depend on their capacity to adapt to the rapidly changing environment.

Here, we document the parallel acceleration of biological aging, a biomarker of cumulative lifetime exposure to stressors, and the erosion of fitness across more than a half-century of recent climate change in an intensively studied polar bear (*Ursus maritimus*) population (Fig. 1). Given that the planet will likely continue to warm for several centuries regardless of near-term emission reductions(*6*), we then explore the capacity of polar bears in our study population to adapt to future climate change by estimating adaptive potential (Figure 1). The Arctic has warmed approximately four times faster than the global mean and is now 3°C warmer, on average, than at the onset of rapid warming(*7*). If current emission targets are met, the rest of the planet will have warmed by ∼3°C by the end of the century(*8*). Our findings thus provide insight into organismal decline due to climate change at the cellular level and whether we might expect widespread adaptive change by populations to levels of warming that will soon be reached globally, particularly for species adapted to narrow climatic conditions.

**Fig. 1.**
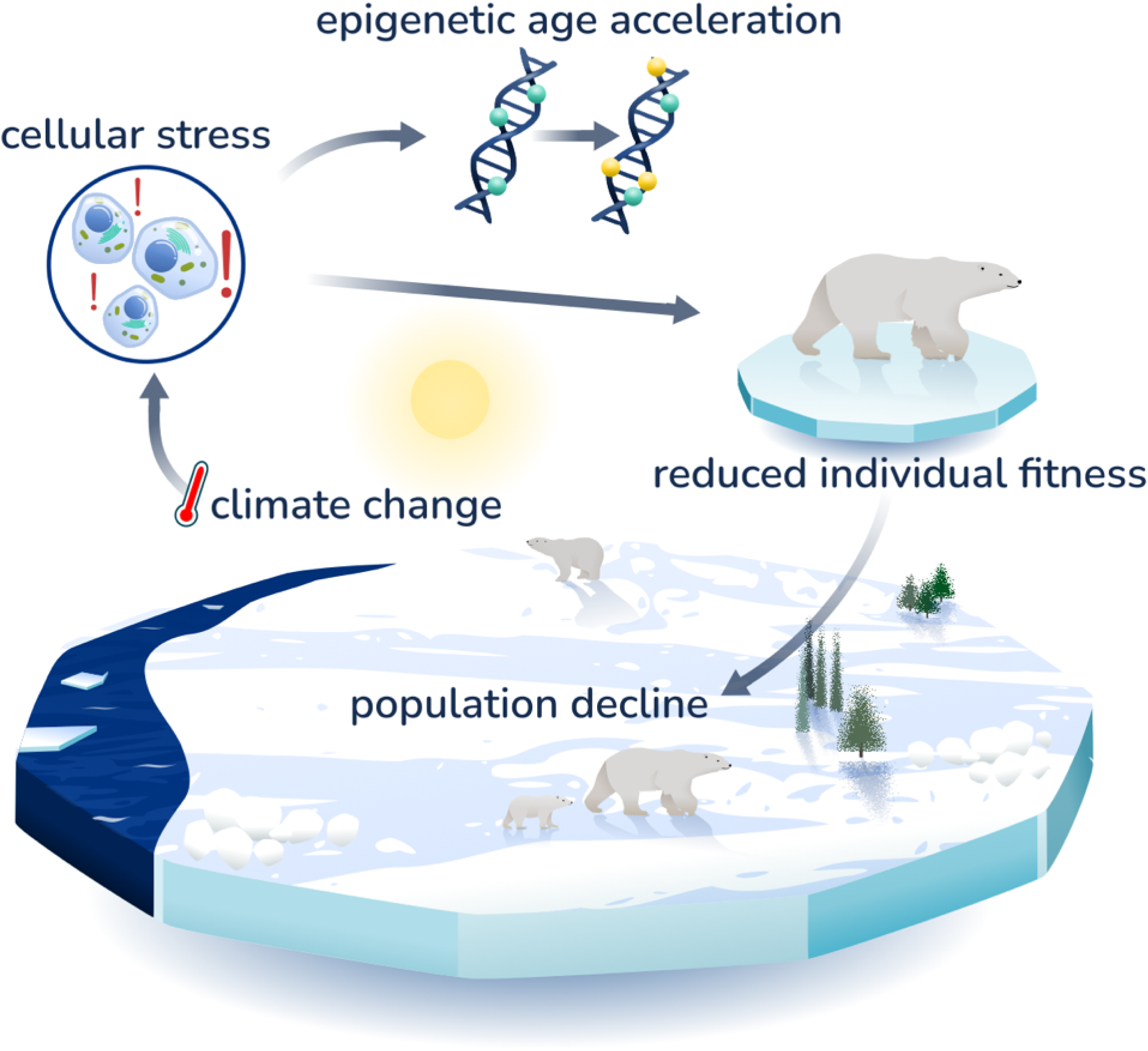
Epigenetic age acceleration from cellular stress parallels reduced fitness, limited adaptive potential, and population decline in polar bears under climate change. Global concerns over declining polar bear populations prompted the formation of an international conservation agreement for the species in the mid-1960s, initiating annual sampling and individual-based monitoring of polar bears in western Hudson Bay. Using the long-term individual-based dataset, we explored how climate warming impacted individuals and populations. Climate change increased the lifetime exposure of individuals to stress, detectable in epigenetic markers from archived tissue samples. This stress also compromised fitness, reducing adaptive potential and driving population declines.

The western Hudson Bay polar bear population is at the southern edge of the species range and has been subject to standardized annual sampling and individual-based monitoring since 1980. Polar bears rely on sea ice for travel, mating, and as a platform to hunt their primary prey, ringed (*Pusa hispida*) and bearded (*Erignathus barbatus*) seals. After sea ice retreats in spring, their prey becomes inaccessible, and polar bears must fast, relying on accumulated fat reserves for growth, reproduction, and survival(*9*). As a result of air and sea surface temperature anomalies associated with warming(*10*), the ice-free season in the Hudson Bay region has lengthened by approximately ten days per decade since the early 1980s(*11*, *12*). During this time, the western Hudson Bay polar bear population declined by 35%(*11*). Sea ice loss is firmly linked to this decline and is the most significant climate-related threat to this population(*11–15*). Longer ice-free seasons increase the bears’ fasting period on land, and each additional day of fasting requires metabolizing approximately one kilogram of body mass(*16*). Bears also risk losing stored body mass as thinning winter ice and rapid spring melts force longer swims between ice floes. Swimming is five times more energetically expensive for bears than walking, and dramatically longer swims are required for even small changes in sea ice(*17*).

### Accelerated aging with climate warming

In 1956, Hans Selye observed that ‘Every stress leaves an indelible scar, and the organism pays for its survival after a stressful situation by becoming a little older’(*18*). This notion predicts that exposure to stress across lifetimes should increase biological aging rates, which should, in turn, reduce fitness. Biomedical research has since established that an organism’s cumulative experience of stressful environmental conditions is indeed reflected in its biological age. When an organism experiences excess stress, molecular wear and tear make its cells biologically older than their chronological age suggests. This phenomenon, known as biological age acceleration, is associated with age-related declines in health and well-being in humans and lab animals(*19*, *20*). If climate warming accelerates the onset of age-related phenotypes through cumulative lifetime stress in wild populations, fitness declines should follow, affecting their capacity to adapt to changing environments.

We measured biological age acceleration in western Hudson Bay polar bears using an epigenetic approach recently developed in biomedicine(*21–23*). Epigenetic age acceleration is the residual difference from the linear relationship between chronological and epigenetic age(*24*). This makes it a consistent estimate of biological age acceleration across chronological ages. Faster epigenetic aging is associated with the expression of age-related phenotypes(*25*), but understanding causal links between epigenetic aging and the aging process is an active area of research(*26*). For example, epigenetic aging has been associated with some of the classic hallmarks of aging, such as deteriorating mitochondrial function, nutrient sensing, and stem cell composition, but not with other aspects of aging, such as cellular senescence, telomere shortening, or genomic stability(*27–29*). Regardless of the cellular mechanisms underlying epigenetic age acceleration, it is a reliable biomedical measure of cumulative experiences of environmental stressors across lifetimes(*30–32*), increasing, for example, in humans in response to smoking(*21*), early life adversity(*33*), and declining physical health(*22*). Epigenetic age acceleration is also one of the best biomarkers of morbidity and all-cause mortality in humans(*34*, *35*). In non-human wild animals, epigenetic age acceleration slows during hibernation(*36*) and has been linked to individual rank in social groups(*37*).

Epigenetic age is measured through DNA methylation, a process in which methyl groups are bound to DNA at cytosine guanine dinucleotides (CpG or CG sites)(*19*). This process plays a role in cell fate and gene regulation(*38*). DNA methylation patterns at some CpG sites change so predictably across lifespans that they can be used to build “epigenetic clocks” for predicting chronological age in humans(*24*), mice(*39*), and many other mammals(*40–45*). Epigenetic clocks predict chronological age two- to three-fold more accurately than earlier biological aging approaches such as telomere shortening(*46*). Critical for our purposes, epigenetic age acceleration can be estimated from archived tissue sampled from individuals with known chronological ages(*24*). Archived samples are available for the western Hudson Bay population dating back to the 1980s, providing a lens through which we could view changes in the accumulation of stress across lifetimes in polar bears, starting before the onset of significant climate warming through the recent period of rapid warming.

We built an epigenetic clock for polar bears with archived tissue samples (Methods – *Building the epigenetic clock for polar bears*). Our clock (Data S1) is based on blood and skin tissue DNA methylation measurements from 144 male and female individuals evenly sampled across ages 0–30 between 1988–2016 (Data S2; Methods – *Field data collection*). We used epigenome-wide association surveys followed by elastic net regression to identify 125 CpG sites whose values can be used to calculate epigenetic age in polar bears. We narrowed these sites down from an initial 33,674 candidate CpG sites from the mammalian DNA methylation array of highly conserved regions(*47*) that align to the polar bear genome (Data S3). The selected sites were strongly associated with chronological age but unrelated to sex or tissue type (Fig. S1).

Our clock (Fig. 2A) estimated chronological age in an independent validation set of 228 samples from 162 individuals not used for clock development with a Pearson’s correlation of 0.94 (5^th^ to 95^th^ percentile of bootstrap-sampled clocks 0.92–0.95) and a median absolute error of 2 years (5^th^ to 95^th^ percentile of bootstrap-sampled clocks 1.74–2.44), 7–10% of the typical polar bear lifespan(*48*). This performance is on par with the most accurate epigenetic clocks built for humans and other wildlife(*24*, *41*, *42*). Our polar bear clock also estimated chronological age more accurately and consistently across blood and skin samples than the recently developed universal pan-mammalian clocks(*40*) (Fig. S2). We used our epigenetic clock to estimate epigenetic age acceleration through time for the 162 individuals not used for clock development.

**Fig. 2.**
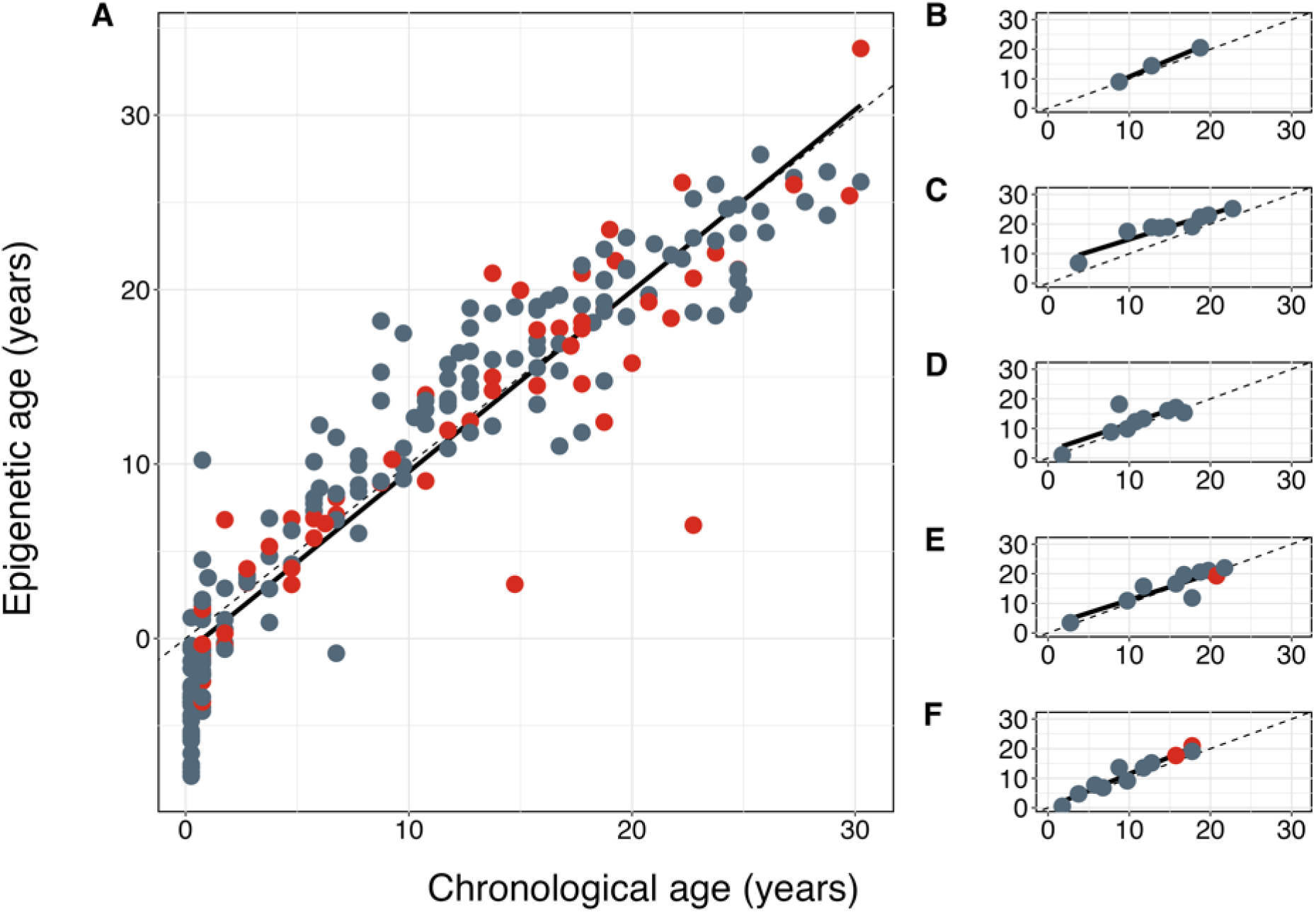
The polar bear epigenetic clock tracks chronological age in the western Hudson Bay population. Epigenetic clocks estimate age using age-related methylation patterns at cytosine guanine dinucleotides on DNA molecules. The clock was developed with samples from n = 144 male and female individuals aged 0–30 sampled between 1988–2018. **(A)** shows n = 228 samples not used for clock development, aged 0–30 and sampled between 1988–2023. **(B–F)** show five individuals sampled repeatedly over their lifetimes. Points represent samples from blood (red) and skin (grey), the dotted lines show a 1:1 relationship between chronological and epigenetic age, and the solid lines are regression lines between epigenetic age and chronological age (median absolute error = 2 years; Pearson’s correlation 0.94). Points above this line indicate age acceleration—samples appearing epigenetically older than their chronological age.

We found evidence for epigenetic age acceleration through time that paralleled climatic warming and lengthening ice-free periods. Polar bears born more recently aged faster epigenetically as the climate warmed (Table 1; Fig. 3). Both males and females responded similarly (Table S1, Fig. S3), and the relationship was consistent between blood and skin tissue (Table S1, Fig. S4). On average, bears born in the 2020s were epigenetically 2.7 (interquartile range 1.4) years older than bears born in the 1960s (Fig. 3). This relationship held when we repeatedly re-sampled the training and validation data with replacement, re-fit the clock, and re-analyzed the relationship between birth year and epigenetic age acceleration (Fig. S5). Considering the typical 15–20-year lifespan for polar bears in the wild(*48*), a 2.7-year increase in epigenetic age is equivalent to a 13.5–18% increase in lifetime aging rate.

**Table 1.** Climate change is linked to epigenetic aging and fitness declines in an intensively studied polar bear population in western Hudson Bay, Canada. Bears born more recently and those that reproduce earlier in life age faster epigenetically. The fitness benefit of earlier reproduction, estimated using lifetime reproductive success, declined for later-born bears. We report the coefficients (ß) and 95% credible intervals (CrI) from Bayesian regression models testing relationships between epigenetic age acceleration, year of birth, age at first reproduction, and lifetime reproductive success. The probability of direction (P direction) describes the probability that a coefficient is either positive or negative, expressed as a percentage between 50% and 100%. The Bayesian R^2^ describes the proportion of variance explained by the model. For all models, we used conservative, weakly informative priors with mean = 0 and standard deviation = 1. We fit all models using the brms package v2.20.4 in R v4.3.1, with 4 chains and 10,000 iterations, including 5,000 warmup iterations. Posterior predictive checks are in Fig. S7.

| Variable | Covariates | $\beta$ (95% CrI) | Bayesian $R^2$ | P direction |
| --- | --- | --- | --- | --- |
| Age acceleration<sup>§</sup><br>(n = 162) | Year of birth | 0.21 (0.06, 0.36) | 0.05 | 99.6% |
| Age acceleration<sup>§</sup><br>(n = 76) | Age at first reproduction | -0.28 (-0.50, -0.06) | 0.08 | 99.2% |
| Lifetime reproductive success<sup>†</sup><br>(n = 896) | Age at first reproduction | -0.10 (-0.15, -0.06) | 0.08 | 100% |
|  | Year of birth | -0.17 (-0.22, -0.12) |  | 100% |
|  | Age at first reproduction: Year of birth | 0.04 (0, 0.09) |  | 96.5% |

<sup>†</sup>Bayesian generalized linear models were specified using a negative binomial response distribution with a log link function.
<sup>§</sup>Bayesian linear models were specified using an identity link function.

**Fig. 3.**
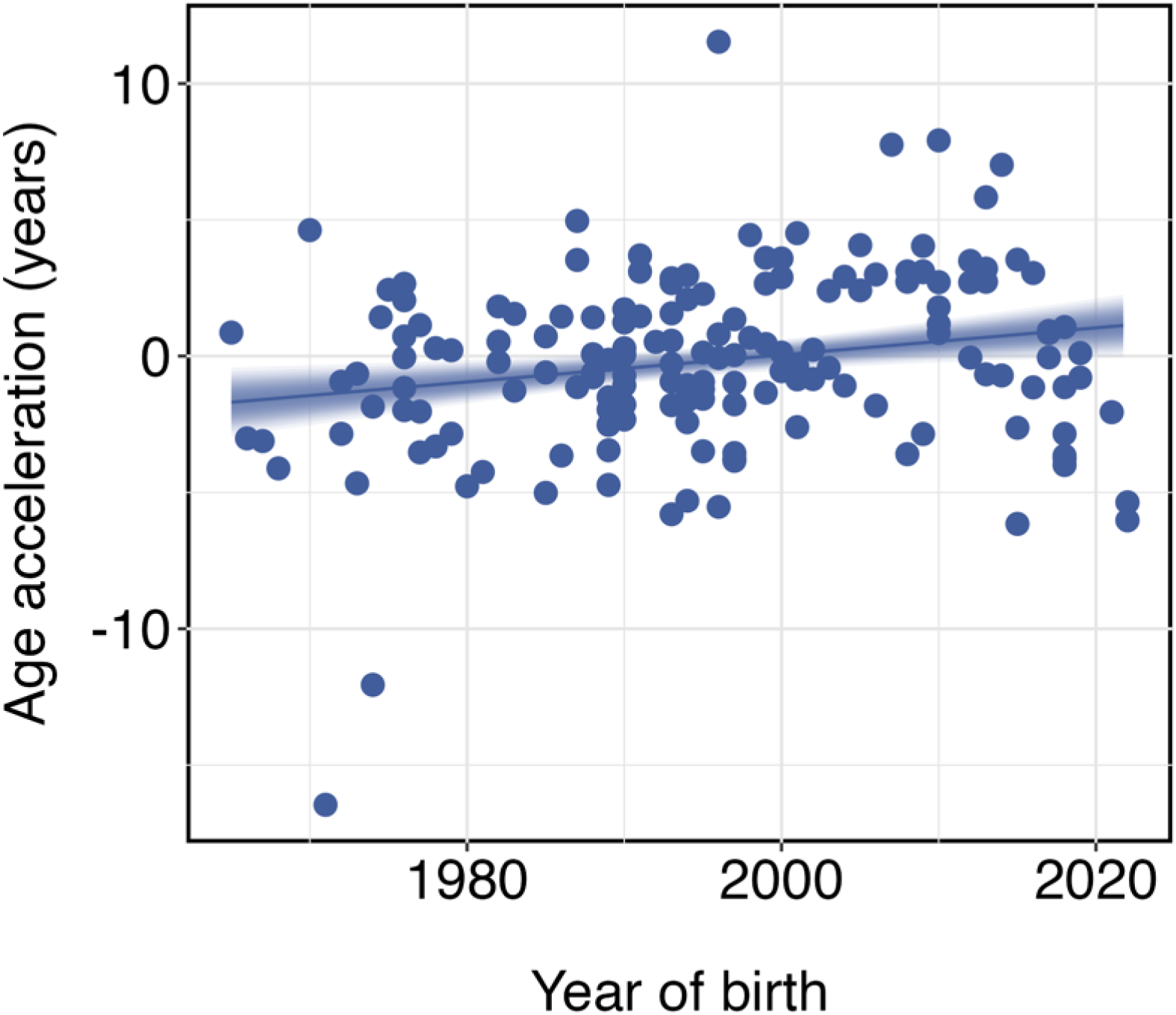
Epigenetic age, a marker of cumulative lifetime stress, has accelerated through time for polar bears in western Hudson Bay, Canada. Points are observed values of epigenetic age acceleration for n = 162 individuals born 1965–2022 and sampled between 1988 and 2023. The line and ribbon are the mean and 95% credible intervals around the posterior distribution of the Bayesian linear model (ß = 0.21, 95% credible interval 0.06, 0.36).

Epigenetic aging rates were consistent across individual lifetimes. Our polar bear clock accurately tracked chronological ages in individuals with repeat samples (Fig. 2B–F). We also tested whether aging rates were being overestimated because of faster epigenetic aging in younger individuals, a phenomenon known to occur in humans(*49*). When we refit the clock and repeated the downstream analysis using only samples from mature bears, we found the same epigenetic age acceleration through time, but the clock fit with only mature bears was less accurate than our original clock (Fig. S6). This loss of accuracy indicates that early-life epigenetic age acceleration is important for understanding age acceleration across the lifespan and that the inclusion of young bears did not adversely bias our clock.

Notably, the increased aging rate we found over time is likely a conservative estimate because of our sampling approach. We evenly selected samples from individuals across age classes (Methods – *Building the epigenetic clock for polar bears*), but polar bear survival declines substantially as they approach their early 20s(*50*). In humans, epigenetic age acceleration is associated with increased morbidity and mortality(*22*, *25*). If this association holds for polar bears, the 20–30-year-old bears available to sample were likely among the healthiest of their cohorts, meaning we likely oversampled healthy older bears and might have underestimated epigenetic age acceleration.

### Evolutionary change and adaptive potential

We next sought to link our estimates of epigenetic age acceleration to predictions about aging rates drawn from well-established life history theory that predicts trade-offs between lifespan and investing in reproduction(*51*). Increased biological aging rates through time should eventually be associated with the erosion of fitness and the loss of adaptive potential as individuals express age-related phenotypes associated with reduced fitness earlier in life(*51*, *52*). We thus also tested for fitness benefits associated with reproducing early in life and their change over time— reproducing early in life should enhance lifetime reproductive success in uncertain environments where survival and reproduction later in life is uncertain. Finally, we estimated adaptive potential given we are now certain of future environmental change.

We first explored relationships between epigenetic age acceleration and the timing of first reproduction. Individuals take energy and nutrients from the environment and allocate them to self-maintenance and reproduction. Resources devoted to one area are not immediately available for the other, meaning the optimal allocation of energy to self-maintenance and reproduction depends on the ecological setting(*51*). Because reproduction is energetically expensive, classic life history theory predicts reproducing early will come at the expense of self-maintenance and longevity. This trade-off predicts that individuals who reproduce early in life should age faster epigenetically than those who first reproduce later. Indeed, we observed a negative relationship between mean epigenetic age acceleration and age at first reproduction for 76 bears with both estimates of age acceleration and known ages at first reproduction (Table 1; Fig. 4). This finding, predicted by well-established theory(*51*), suggests that epigenetic age acceleration reflects a biologically meaningful age-associated component of polar bear life histories and the energetic trade-off between the cost of early breeding and longevity.

**Fig. 4.**
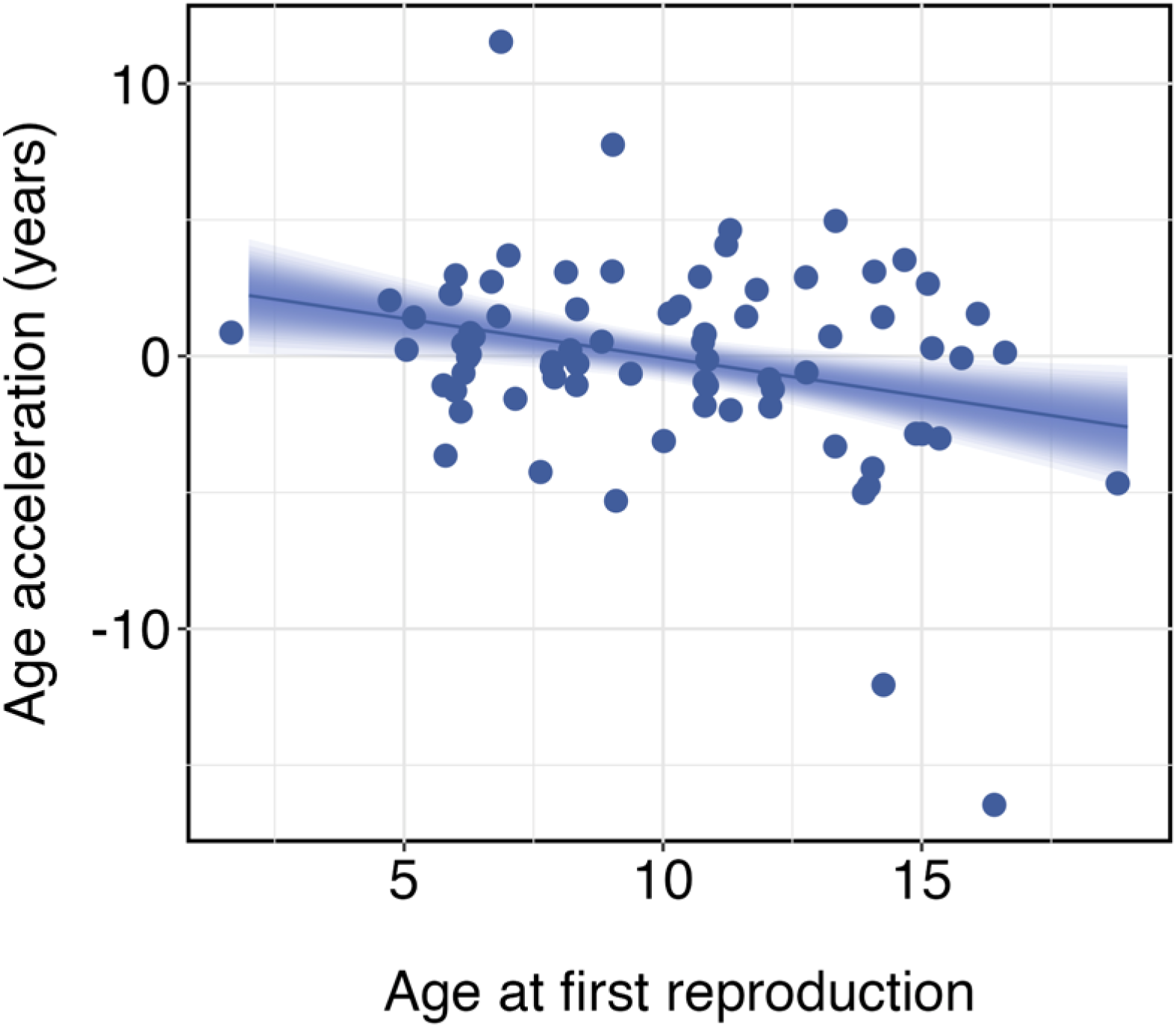
Polar bears that first reproduced at younger ages aged faster epigenetically. Points represent observed values of epigenetic age acceleration from n = 76 individuals born 1965– 2010 and sampled between 1988–2016 for which age at first reproduction is known. The line and ribbon represent the mean and 95% percentile interval around the posterior distribution estimated using a Bayesian linear model (ß = -0.28, 95% credible interval -0.50, -0.06).

Life history theory also predicts that when environments are harsh, and survival and reproduction late in life are uncertain, we should see fitness benefits associated with investing energy in reproducing earlier in life, even if early reproduction accelerates aging(*52*). To explore this possibility, we tested for fitness benefits from breeding early in life(*51*, *52*). We expected early-life reproduction to have fitness benefits for western Hudson Bay polar bears as this population experiences a harsh environment with little certainty in terms of survival and reproduction late in life. We estimated lifetime reproductive success, a common measure of fitness, for 896 individuals from the long-term study (Methods – *Estimating life history traits*). We found that from the 1960s through the 1980s, early reproducing bears had the highest lifetime reproductive success, suggesting that reproducing early was indeed adaptive before the onset of rapid warming (Table 1). However, the fitness advantage of reproducing early in life declined through the 1990s. By 1995, bears produced the same number of offspring over their lifetimes regardless of how young they were when they first reproduced (Fig. 5). The fitness advantage of reproducing early in life has eroded through time in parallel with warming and population-wide epigenetic age acceleration.

**Fig. 5.**
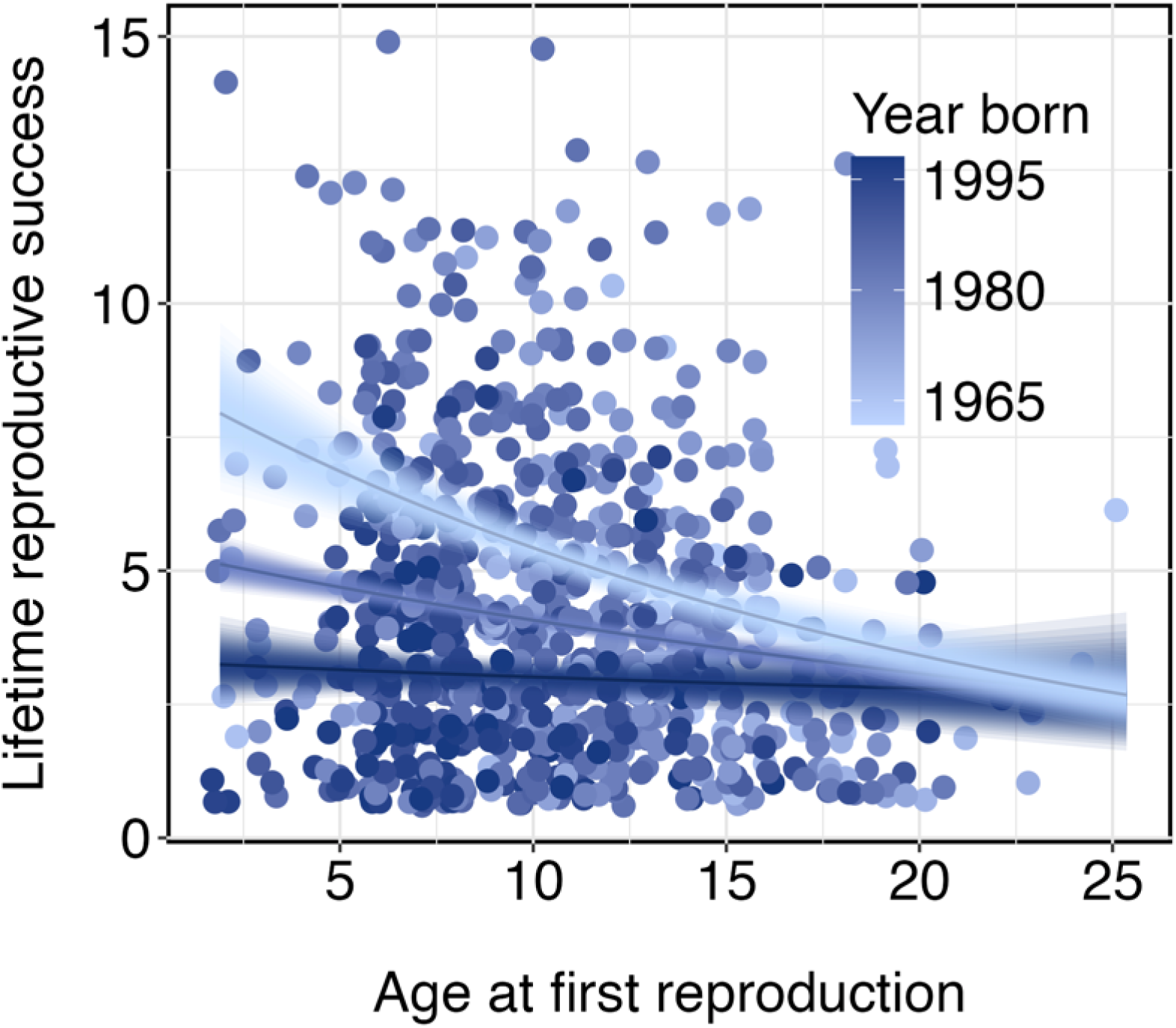
Climate change lowered the fitness benefit of polar bears of early-age reproduction in polar bears. Reproducing at younger ages is energetically costly but confers a net fitness benefit in environments with reproductive uncertainty. Polar bears born in the 1960s in western Hudson Bay had higher lifetime reproductive success when reproducing younger (ß = -0.10, 95% credible interval = -0.15, -0.06). However, this benefit declined for bears born in later decades, disappearing entirely by the mid-1990s (ß = 0.04, 95% credible interval 0, 0.09). Points represent individual observations, and lines and ribbons are the predicted means and 95% credible intervals of the posterior distribution from the Bayesian generalized linear model.

Finally, building on associations we found between epigenetic age acceleration, reduced fitness, and the warming climate over time, we explored the overall rate of recent adaptive evolution in the western Hudson Bay polar bear population and its capacity to adapt to future environmental change. All credible climate forecasts project ongoing warming for the foreseeable future. This means the population’s persistence will depend considerably on its capacity to adapt genetically to the changing environment. We assessed the population’s adaptive potential by estimating the additive genetic variance in individual fitness, which can be thought of as the heritable genetic variation underlying the ability of individuals to reproduce(*53–55*). This approach cannot accurately estimate the contributions of genetic variation related to dominance and epistasis; however, most genetic variation underlying polygenic traits is additive(*56*).

Using our estimates of lifetime reproductive success and relatedness from the population’s 4,634 individual pedigree (923 dams, 443 sires) that spanned the 60-year study(*57*), we fit an animal model(*58*) and found the additive genetic variance underlying lifetime reproductive success in this population is approximately zero (*V*_A_ (w) = 0.008; Table S2). This suggests that selection on traits related to individual fitness is not contributing to adaptive change in this population and that variation in lifetime reproductive success is driven by the environment. This lack of evidence for adaptive change is consistent with observed low survival rates and the population’s recent decline(*59*). Many factors, including gene flow, genetic drift, mutations, and changing environments, regularly place upper limits on adaptive evolution in natural populations(*53*). The magnitude and pace of climate warming appear to limit any adaptive evolution in the western Hudson Bay polar bear population.

### Conclusions

We found parallel increases in epigenetic age acceleration in polar bears and the erosion of their fitness through time associated with climate warming. Epigenetic age acceleration is detectable at the cellular level and accumulates over time(*24*), making it well-suited for estimating lifetime stress associated with climate warming. We found that despite some warmer years, epigenetic age acceleration increased across polar bear lifetimes following local warming trends. As predicted for a population experiencing accelerated aging and fitness declines, current rates of environmental change appear to be outpacing adaptive evolution. With prolonged exposure to harsh environments, abundance declines(*11*), and little evidence for adaptive capacity, the western Hudson Bay polar bear population faces an uncertain future.

Our findings of accelerated aging, the erosion of fitness with warming, and limited evidence for adaptive potential given current environmental conditions are instructive for understanding whether and how other populations might respond adaptively to environmental change in the coming decades. Warming in the Arctic has abruptly altered the ecosystem, and this abruptness has outpaced the adaptive capacity of western Hudson Bay polar bears. Recent modelling suggests that similarly abrupt exposures to intolerable temperatures across large portions of species’ ranges could be widespread in the coming decades(*60*). At the current rate of warming, more than 30% of species could be exposed to temperatures beyond those they evolved to tolerate by 2100(*60*, *61*). As species pass their thermal thresholds, the capacity for populations and ecosystems to adapt will likely diminish.

With the recent signing of the United Nations Montreal-Kunming Global Biodiversity Framework, monitoring and conserving adaptive potential has become mandated in international policy. This is a major positive step toward safeguarding biodiversity. Direct estimates of adaptive potential similar to ours require long-term individual-based studies of wild populations, which are very valuable and rare, particularly in polar regions, due to logistical constraints and costs associated with maintaining long-term research. Our results, and forecasts of future species exposure to warming(*60*, *61*), warn that adaptive responses to warming could be the exception rather than the rule. Significant conservation efforts on all fronts and concerted efforts to halt warming will be necessary to safeguard biodiversity for future generations.

## Materials and Methods

### Summary

We studied biological aging, fitness, and adaptive potential in the western Hudson Bay polar bear (*Ursus maritimus*) population. Using a pedigree previously constructed for the population(*57*), we estimated ages at first reproduction and lifetime reproductive success. We measured biological aging by first building an epigenetic clock for polar bears and then using it in independent samples to gauge the rate at which the cells of individual bears from the population aged epigenetically relative to their chronological age. Using regression models, we tested for an increase in the rate of epigenetic aging over time, relationships between age at first reproduction and the rate of epigenetic aging, and changes in lifetime reproductive success with age at first reproduction over time. Finally, we used an animal model, a type of mixed-effects model that estimates the additive genetic variance of traits like lifetime reproductive success(*2*), to estimate the capacity of western Hudson Bay polar bears to adapt to climate change.

### Field data collection

Since 1966, polar bears have been captured in northeastern Manitoba near Churchill, Canada, as part of a long-term study(*57*). Bears are chemically immobilized, sexed, and fitted with unique ear tags and tattoos on the upper lip for later identification in case of recaptures. Skin samples are extracted either from pinna tissue remaining after ear-tagging or using a biopsy punch of superficial rump fat(*57*). Blood samples are drawn from femoral blood into a sterile Vacutainer and stored at -80° C(*57*). All capture and handling protocols are approved annually by Environment and Climate Change Canada’s Prairie and Northern Region Animal Care Committee and wildlife research permits are issued by the Province of Manitoba and by Parks Canada. A standardized sampling program was initiated in 1980 and continues today, with the exclusion of 1985 and 1986. As part of this program, adult females and their cubs of the year are sampled in February and March. Chronological age is either derived from known years of birth for cubs-of-the-year or based on cementum annulus deposition from an extracted vestigial premolar tooth for bears first captured as adults(*62*).

### Estimating life history traits

We estimated polar bear life history traits using previously established pedigree relationships. The western Hudson Bay polar bear population pedigree(*57*) contains 4,634 polar bears (443 sires, 923 dams, and 1,130 founders, i.e., individuals of unknown parentage) from over six generations sampled between 1966 and 2016. Field sampling data from females and cubs-of-the-year provided offspring-dam associations. Additional linkage information came from parentage analyses using multi-locus microsatellites to genotype individuals(*57*). We removed three individuals whose sex classifications were inconsistent with parentage data; these individuals were classified either as male dams or female sires. Additional information about the pedigree construction, capture, handling, and sampling protocols for the western Hudson Bay subpopulation is described in more detail in previous work(*57*).

We used the pedigree to estimate lifetime reproductive success and age at first reproduction for all individuals who were confirmed parents of at least one other bear in the pedigree. We defined lifetime reproductive success as the total number of other bears in the pedigree for which an individual was a confirmed parent. We defined age at first reproduction as the age of the individual when its first known offspring was born. In our analyses considering lifetime reproductive success, we removed individuals with potentially biased data, using only data from individuals born in 1996 or earlier (n = 896 bears). We selected this threshold because any bears born after 1996 would not have reached 20 years—the approximate age of senescence for western Hudson Bay polar bears(*63*)— by 2016, the year the pedigree was completed for the population(*1*). As a result, the late-life offspring of these bears might have been missed in the pedigree, potentially resulting in biased estimates of lifetime reproductive success. While we expected some error in lifetime reproductive success because of gaps in the pedigree, we assumed relative comparisons among individuals were unbiased up to 1996.

### Building the epigenetic clock for polar bears

Epigenetic clocks predict chronological ages based on methylation of CpG dinucleotides, where a cytosine is followed by a guanine(*24*). Many CpGs change with chronological age. The Illumina HorvathMammalMethylChip40 array (Illumina Inc., San Diego, CA, USA; hereafter mammalian array) was designed to analyze methylation at CpG sequences highly conserved across all mammal species, measuring a total of 37,449 unique sequences per sample at a single nucleotide resolution. The high-throughput array can process 96 samples simultaneously, making this approach useful for aging samples from long-term ecological projects that store many samples over multiple years.

To build our epigenetic clock, in 2022, we randomly selected 288 samples from 6,135 blood and skin samples collected from western Hudson Bay polar bears aged 0–30 born from 1965–2018. We stratified sampling based on individual age, tissue type, sex, and year of sample collection. We ensured some samples came from bears sampled more than once over their lifetimes to test for consistency in individual aging rates over time (Figure 2). To test for consistency in DNA methylation rates between tissues, we also included several samples with blood and skin collected simultaneously from the same individual. Our samples included 150 female and 138 male samples from 223 unique individuals, of which 111 were blood and 177 were skin. Five individuals were sampled between 4 and 13 times over their lives, and another 25 individuals were sampled twice. In 2024, we added 96 additional samples from individuals sampled one time each, as recently as 2023, to extend the time frame and sample size covered. We used these additional samples for clock validation but not development. Details of all final samples used for clock development, validation, and downstream analysis are available in Data S2.

We isolated genomic DNA from blood and skin samples using the Qiagen DNeasy Blood and Tissue Kit 250 (Qiagen, Hilden, Germany). We dissected approximately 25 mg of frozen skin samples on a pre-chilled plate placed on dry ice to prevent thawing of the entire tissue. We then cut the skin tissue into smaller pieces, placed it in 1.5 mL microcentrifuge tubes, and digested it overnight in Buffer ATL (Qiagen) with Proteinase K (Qiagen) solution at 56 °C. We also digested 50 µL volumes of blood samples in Proteinase K (Qiagen) and PBS (pH 7.4, 1X, Gibco) solution at 56 °C for 10 minutes. After tissue digestion, we extracted genomic DNA from the samples as per the manufacturer’s protocol (Document #HB-0540-002, Version #04/2016) and eluted the samples in two 100 µL volumes of elution buffer (Qiagen) consecutively to increase yield. We measured the concentration of gDNA using the NanoDrop2000 spectrophotometer (Thermo Scientific, Wilmington, USA). Next, we treated 750 ng of each genomic DNA sample with sodium bisulfite using the EZ-96 DNA Methylation-Gold Kit (shallow-well format) (Zymo Research, CA, USA) as per the manufacturer’s protocol (Document #D5007, Version #2.1.6). We eluted the bisulfite-converted DNA in 12 µL of elution buffer (Zymo Research), after which we amplified 4 µL from each sample to be hybridized onto the mammalian array following the Infinium HD Methylation Assay protocol (Document #15019519, Version #07).

We measured DNA methylation by imaging the hybridized chips on the same day they were stained using the iScan instrument (Illumina Inc., San Diego, CA, USA). We normalized the raw intensity data (IDAT) files from the chip scans using the recommended pipeline in the *minfi* package(*64*) in Rv4.3.1(*65*). The code we wrote for pre-processing is available at https://github.com/ljnewediuk/PB_life-history. Normalized intensity data, hereafter 11 values, quantify the degree of methylation at each of the 37,449 sites on the mammal chip with a value between 0 for no methylation and 1 for 100% methylation at each site.

The design of the mammalian array, while appropriate for any mammal species, presents some practical challenges in terms of accurately quantifying methylation in the genomes of specific species. First, all 37,449 CpG sequences on the array might not bind to the genomes of all species because of species-specific CpG differences. Alternatively, a single sequence might bind multiple times in the genome of a given species because probes were designed with up to three degenerate bases to facilitate matching in case of cross-species mutations(*47*). Sequences that bind multiple times can confound methylation signals coming from multiple sites at once(*47*). Methylation can also vary by sex if CpG sequences are located on the sex chromosomes(*66*), a particularly important concern for the mammalian array because of species-specific locations of CpG sequences on chromosomes.

To minimize potential confounds from sex-specific site methylation and non-binding or multiple-binding probes, we narrowed our probe search space before building our clock. First, we aligned the CpG sequences on the mammalian array to a reference polar bear genome (NCBI Genome assembly ASM1731132v1 https://www.ncbi.nlm.nih.gov/datasets/genome/GCF_017311325.1/) using the *QuasR* package v1.40.1(*67*) in R. We selected only the 33,674 sites that bound uniquely. We also limited our search space with an epigenome-wide association study (EWAS), a technique which correlates phenotypic traits with DNA methylation. We used our EWAS to isolate sites with methylation patterns related to age but not different between sexes. We fit three linear models with site-specific CpG methylation proportions as the response and combinations of age and sex as predictors using the *limma* package v3.56.2(*68*) in R. In the first model, we tested the effects of sex on methylation while controlling for ages and tissue types of samples. We also fit two models including only either blood or skin samples to isolate the effects of age on the proportion of methylation in either tissue. We excluded 3,740 CpG sequences significantly associated with sex (p < 0.05) and 29,573 that were not sufficiently associated with age (p > 10e^-6^). We erroneously excluded another 23 (0.007%) CpG sites. We used a final 3,328 of the 37,449 CpG sites on the HorvathMammalMethylChip40 array to build our clock (Data S3).

We first screened unreliable samples from the data. We screened 12 samples because their scans failed on the iScan, because the samples clustered away from others in a principal components analysis of 11 values, or because the 11 values had high detection p-values, a quality control metric indicating poor discrimination of samples from background values. We plotted the detection p-values on the sample year to rule out any visual pattern indicating the possible deterioration of sample quality over time (Fig. S8). Because we included bears from a single population, we were also concerned that potential relatedness between individuals used to build the clock might bias its predictions. We used the *GeneAlEx 6.5* software(*69*) to assess relatedness between individuals using 24 microsatellites (Data S4). We removed any individuals from the training data with a relatedness index > 0.25. We also removed any individuals with repeat samples from the training data.

We fit our DNA methylation clock (Data S1) using a training set of samples from 144 unique individuals balanced across age, sex, and tissue type from the 278 of our initial 288 samples that passed quality control. We fit the 11 ∼ age clock model using the cv.glmnet function in the *glmnet* package v4.1-8(*70*) in R, setting 11 = 0.5 to combine the benefits of both ridge and lasso regression(*24*). This compromise reduces the variance in age predictions at the cost of some bias. We used 10-fold cross-validation to select the optimal regularization parameter(*24*). We validated our clock by using it to predict the chronological ages of all remaining 228 samples (hereafter the validation dataset), including the remaining n = 134 samples from the original 278 selected in 2022 that we did not use for clock development and 94 of the 96 samples added in 2024 that passed quality control (Data S2). CpG sites and 11 values for the clock are available in Data S1.

We also predicted epigenetic ages in our samples using the universal clock for mammals(*40*) to ensure the predictions from this clock roughly matched our expectations of a linear relationship between chronological and epigenetic age (Extended Data Figures 2 & 3). Lu et al.(*40*) built two pan-tissue mammalian clocks, universal clock 2 and universal clock 3, trained on DNA methylation of 59 tissue types from 185 species and measured using the same HorvathMammalMethylChip40 array we used for our samples. The clocks include transformations to account for age at sexual maturity, gestation time, and maximum lifespan, allowing them to predict the age of any eutherian mammal species with reasonable accuracy(*40*). We aged all polar bear samples (n = 372) using universal clocks 2 and 3, excluding the 12 poor-quality samples that did not pass quality control.

### Statistical analysis—Bayesian regression models

We used Bayesian linear models and Bayesian generalized linear models to test for epigenetic age acceleration over time, for a relationship between age acceleration and age at first reproduction, and a relationship between age at first reproduction and lifetime reproductive success in the validation dataset. We fit all models using the *brms* package v2.20.4(*71*) in R, with four chains and 10,000 iterations, including 5,000 warmup iterations.

For the first two models—age acceleration over time and with age at first reproduction— we used Bayesian linear models, specifying an identity link function and weakly informative prior slopes with a mean of 0 and standard deviation of 1. First, we tested for a relationship between birth year and mean age acceleration in the n = 162 individuals from the validation dataset not used for clock development. We also tested for a relationship between age at first reproduction and mean age acceleration for n = 76 of these individuals with epigenetic age estimates and known offspring.

We next tested for a relationship between age at first reproduction and lifetime reproductive success using a Bayesian generalized linear model. For this model, we specified a negative binomial response distribution with a log link function, using weakly informative prior slopes with a mean of 0 and a standard deviation of 1. We included n = 896 bears from the pedigree born before 1996. We specified an interaction between age at first reproduction and birth year to test how the relationship between age at first reproduction and lifetime reproductive success changed over time.

We performed two additional validations to ensure our epigenetic clock and the relationship we found between birth year and age acceleration were robust. First, we explored the possibility that faster aging in reproductively immature bears biased our estimates of age acceleration. In human populations, children age epigenetically faster than adults(*72*). Climate-related epigenetic age acceleration could have been overestimated across lifetimes if young bears also age faster than adults. To explore this possibility, we created a second clock using the n = 120 samples from our training set from polar bears older than five years, the approximate age of sexual maturity in this species(*73*) (Extended Data Figure 6 A). We then tested this clock on the n = 137 validation samples from bears older than 5 years.

A different set of predictors (CpG sites) might be selected each time an epigenetic clock model is fit (*74–75*). This is because the relationship between methylation changes and epigenetic aging is not considered causal. The mammalian array was designed with sites having conserved age-related methylation patterns, meaning many different combinations of sites might predict age reasonably well.

To ensure our epigenetic clock robustly predicted epigenetic age and estimated the relationship between epigenetic age acceleration and year of birth, regardless of which sites were included in the clock, we repeatedly resampled the clock and repeated the downstream analysis. We randomly split all n = 372 samples into 500 new training and validation sample sets, excluding individuals with multiple samples and siblings from the training set. We then refit the epigenetic clock for each sample set to estimate bootstrap confidence intervals for the clock’s median absolute error and Pearson’s correlation and re-tested the relationship between year of birth and rate of epigenetic aging.

### Statistical analysis—Animal model

We used an animal model(*58*) to estimate the additive genetic variance of lifetime reproductive success for 896 polar bears born before 1996. We used the pedigree data to create a genetic relatedness matrix. We fit this matrix as the random effect ‘*animal*’ to estimate additive genetic variance (*V_A_*). Phenotypic variance (*V_P_*) is partitioned into additive genetic variance (*V_A_*) and a residual variance (*V_R_*) component, which is interpreted as the environmental effect. We further partitioned the residual variance by including maternal variance (*V_M_,* or the identity of individual’s dam) and cohort variance (*V_YBirth_,* or year of birth). We also included sex as a fixed effect in the model. We used a log link function, an inverse-Gamma distribution for the random effect variances, and a wide normal distribution for the prior distribution of fixed effects(*76*). We fit the animal model in the package *MCMCglmm* v2.35(*77*) in R with 1,000,000 iterations and 20,000 warmup iterations. The *MCMCglmm* package allows incomplete pedigrees and uses Bayesian inference and Markov chain Monte Carlo (MCMC) methods.

---

**Fig. S1.**
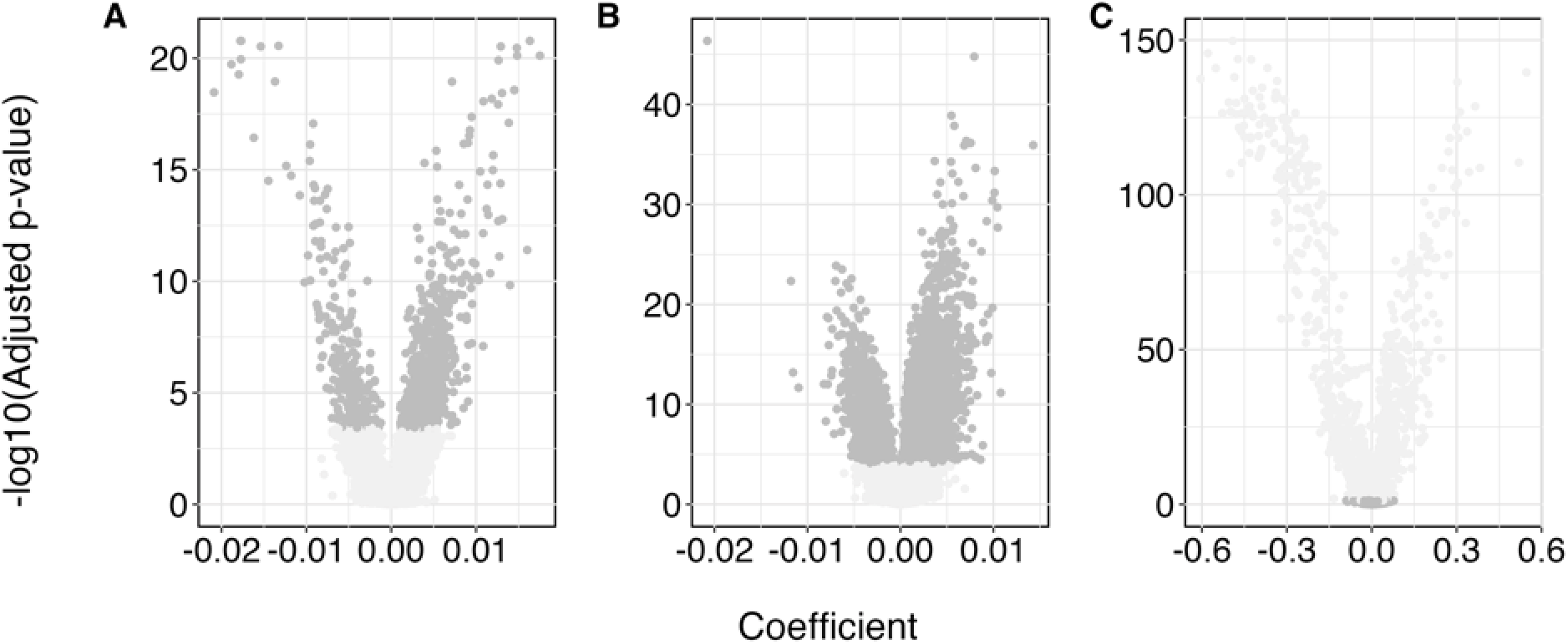
Overview of CpG sites included (dark grey) and excluded from the polar bear clock (light grey). **(A)** Skin sites and **(B)** blood sites were excluded because they were not significantly associated with age (p > 10^-6^), and **(C)** sites from both tissues were excluded because they were significantly associated with sex.

**Fig. S2.**
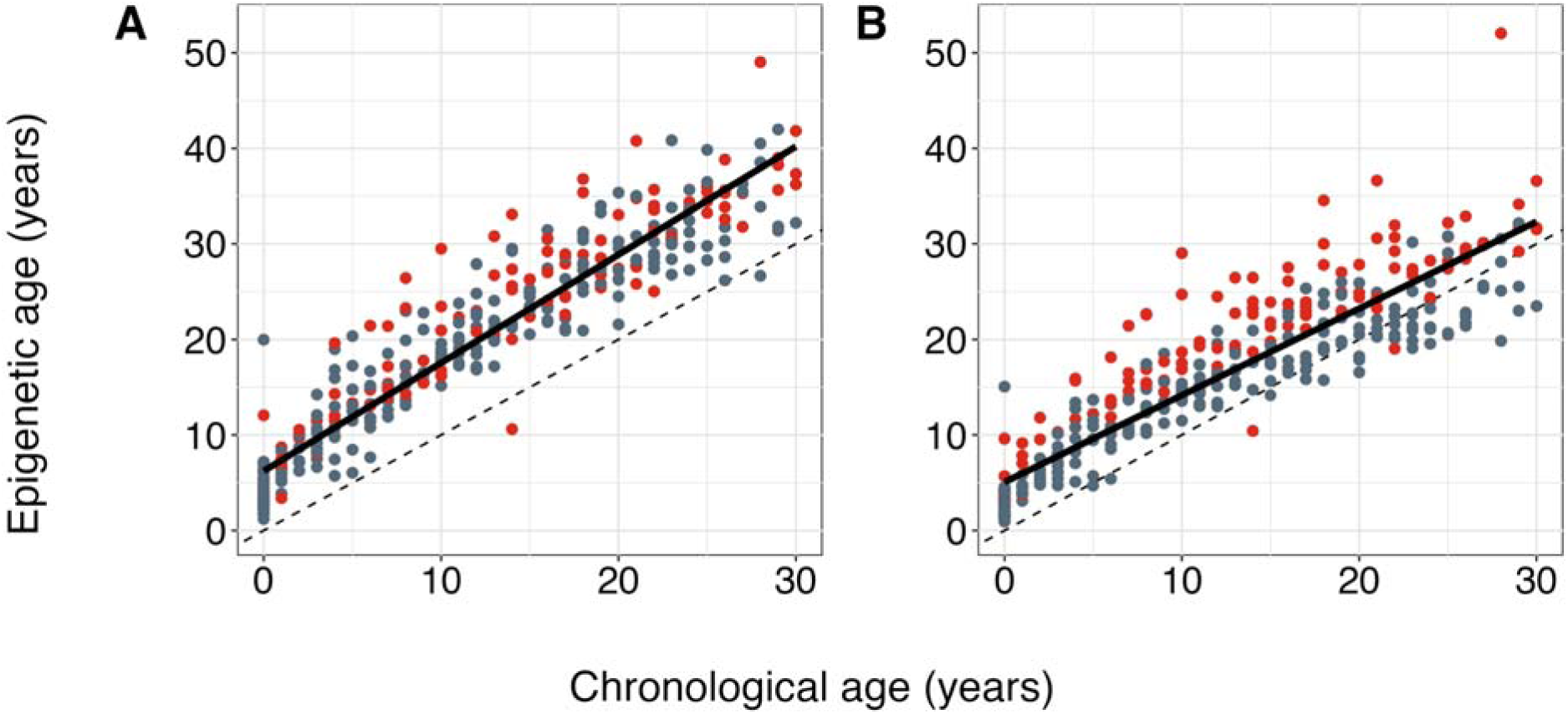
Epigenetic age predictions of n = 372 western Hudson Bay polar bear samples using the universal pan-mammalian clocks(*14*). **(A)** Universal clock 2 (median absolute error = 7.2 years, Pearson’s correlation = 0.94) includes a correction for maximum lifespan (43.8 years) and **(B)** universal clock 3 (median absolute error = 3.8 years, Pearson’s correlation = 0.90 years) includes a correction for age at sexual maturity. Points show individual observations separated by blood (red) and skin (grey) tissue types. The dotted lines show a 1:1 relationship between chronological and epigenetic age, and the solid lines are regression lines between epigenetic age and chronological age. Both clocks overestimate epigenetic ages relative to the polar bear clock (Fig. 2), and universal clock 3’s tissue bias overestimates the ages of blood samples relative to skin.

**Fig. S3.**
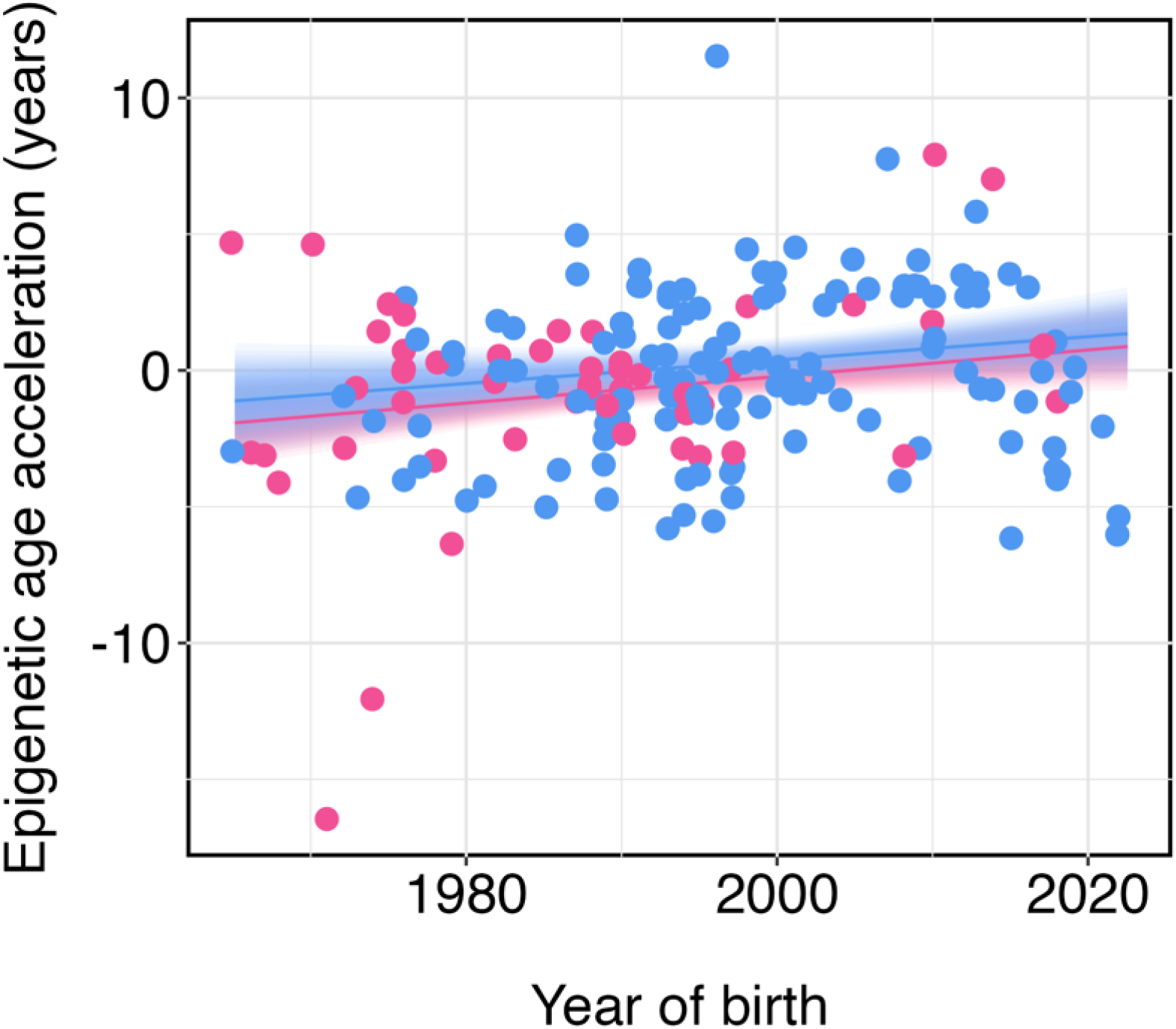
Epigenetic age accelerates with birth year for both male and female polar bears. Points are observed values of epigenetic age acceleration for n = 92 female (pink) and n = 70 male (blue) individuals born 1965–2022 and sampled between 1988 and 2023. The line and ribbon are the mean and 95% credible intervals around the posterior distribution of the Bayesian linear model with an interaction between sex and birth year (model results in Table S1).

**Fig. S4.**
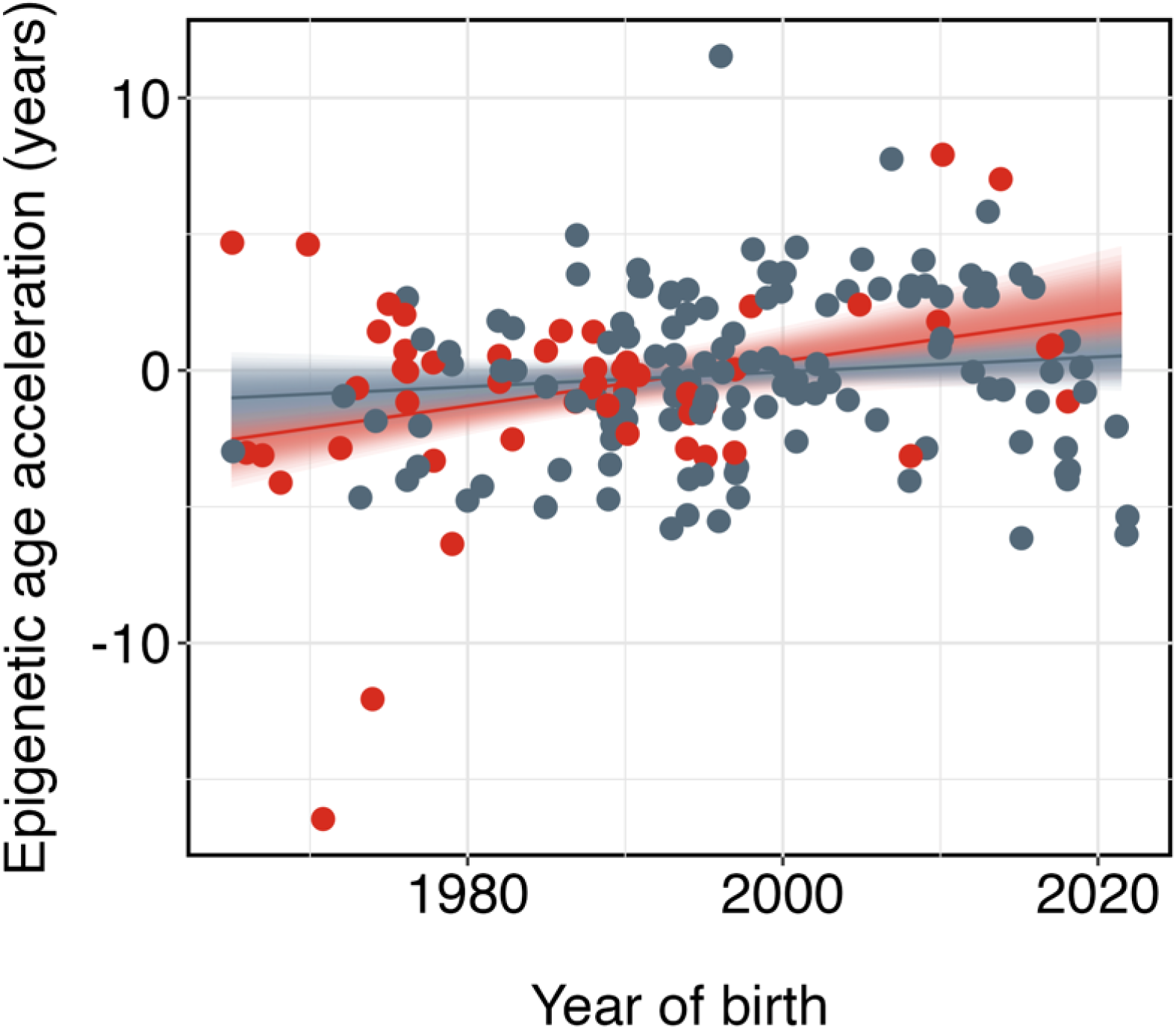
Epigenetic age accelerates with birth year for both tissue types. Points are observed values of epigenetic age acceleration for n = 51 blood (red) and n = 128 skin (grey) tissue samples from individuals born 1965–2022 and sampled between 1988 and 2023. The line and ribbon are the mean and 95% credible intervals around the posterior distribution of the Bayesian linear model with an interaction between tissue type and birth year (model results in Table S1).

**Fig. S5.**
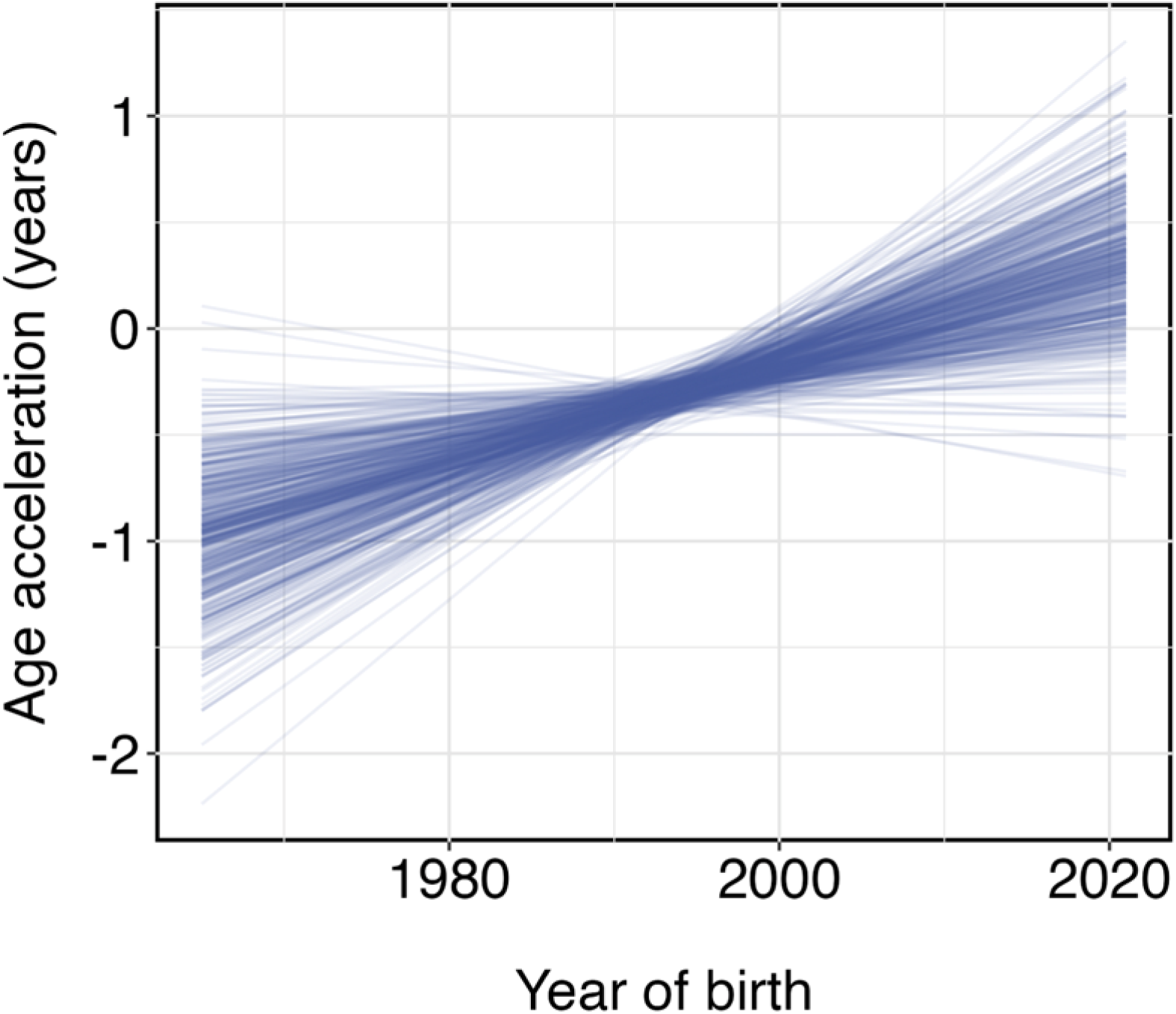
Posterior predictive means from 500 bootstrap-sampled clocks, consistent with our main results. We repeatedly sampled with replacement from 372 male and female polar bear skin and blood samples ages 0–30, born 1965–2022 and sampled from 1988–2023. We used each bootstrap sample to construct a new epigenetic clock. We then used the clock to predict epigenetic age and age acceleration in the remaining individuals not used for clock construction. Each line is the posterior predictive mean from a Bayesian linear model, age acceleration ∼ birth year, fit using the results from each bootstrap sample (mean slope across bootstrap samples = 0.03, 95% credible interval = -0.01, 0.06).

**Fig. S6.**
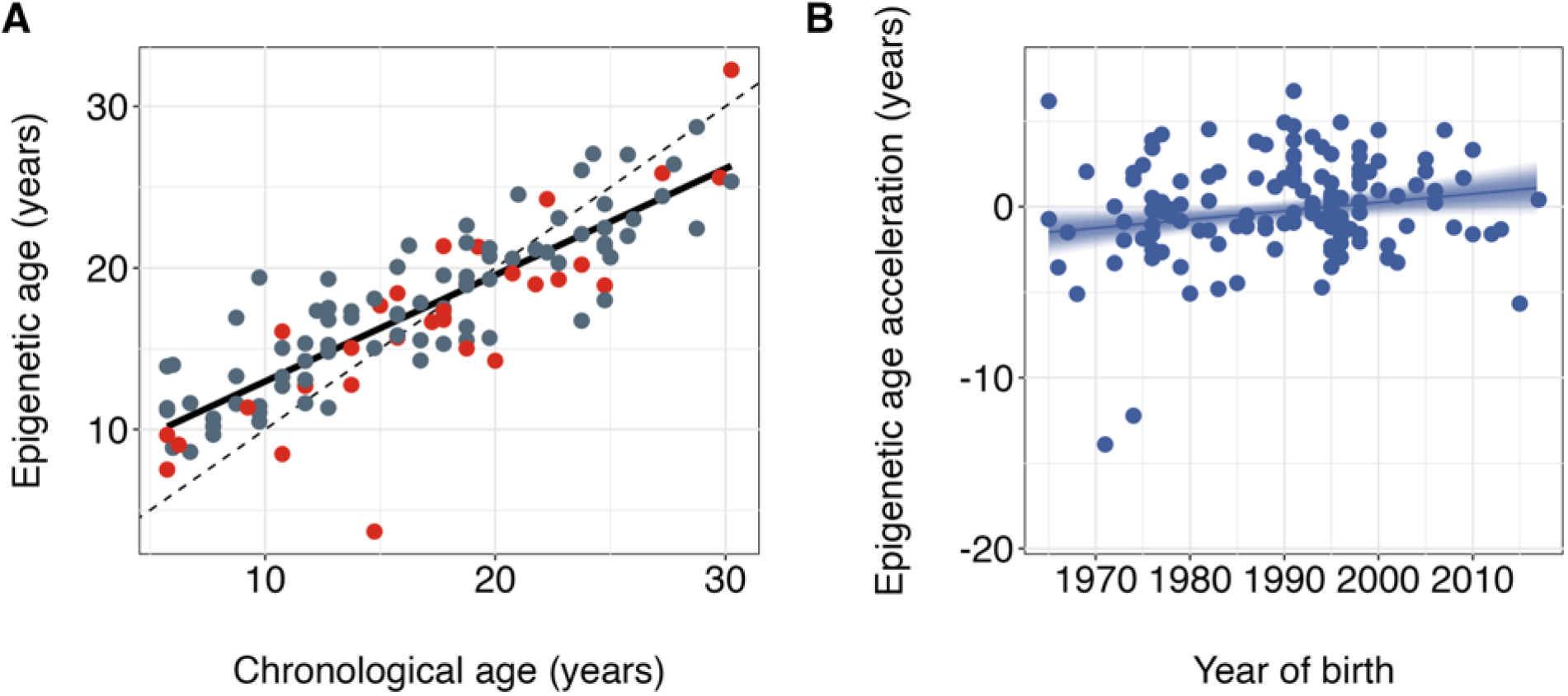
An epigenetic clock constructed with only sexually mature polar bears predicts age less accurately, indicating early-life experiences are important for understanding epigenetic aging rates later in life. **(A)** Epigenetic age predictions for n = 137 blood (red points) and skin (grey points) samples from sexually mature bears aged 5 years and older, predicted using a clock built with n = 120 samples from sexually mature individuals. The clock predicts age with a median absolute error of 2.5 years and Pearson’s correlation of 0.85, which is notably less accurate than the clock that includes young bears (Fig. 2). **(B)** Epigenetic aging rates accelerate over time for 88 bears aged 5 years and older. The line and ribbon represent the mean and 95% credible interval around the posterior distribution estimated using a Bayesian linear model (ß = 0.21, 95% credible interval 0, 0.42).

**Fig. S7.**
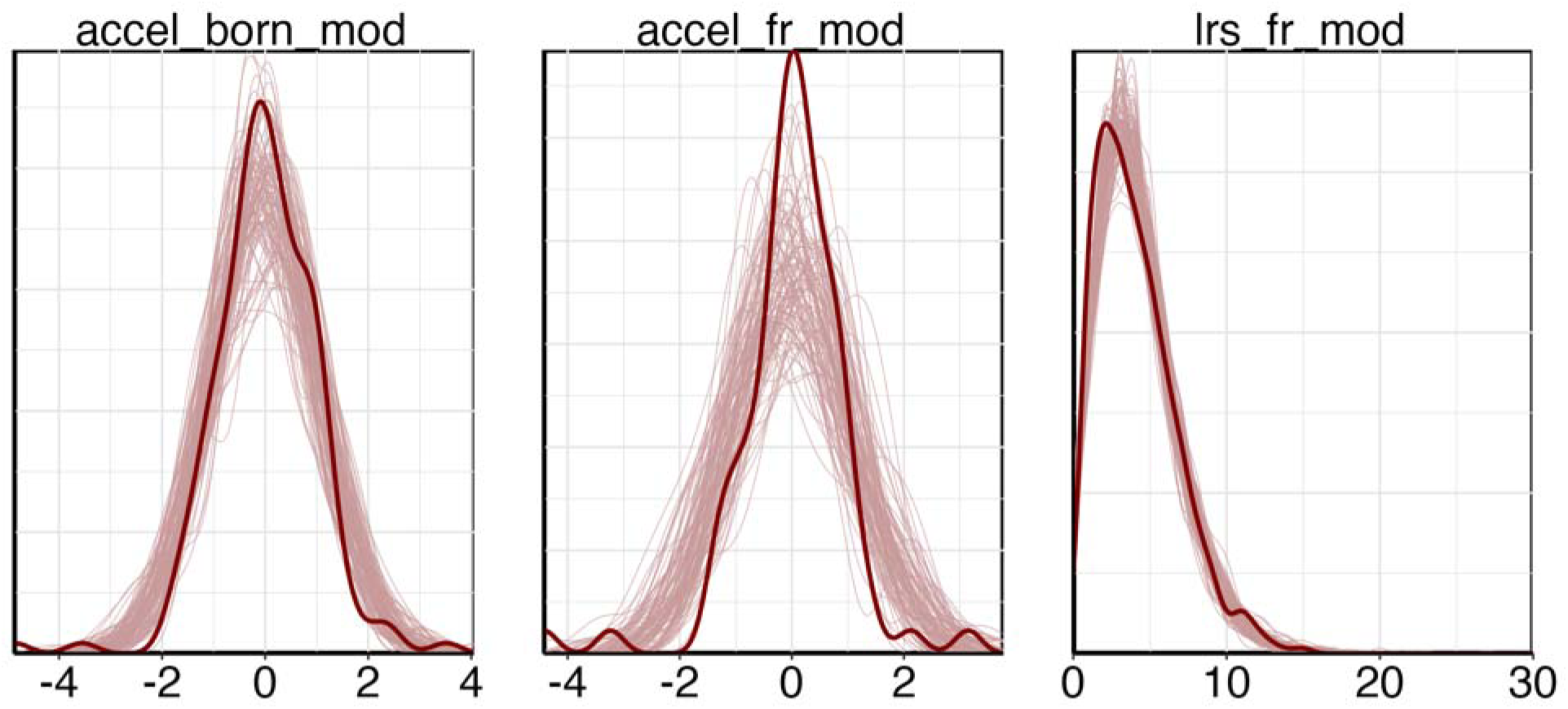
Posterior predictive checks for Bayesian regression models described in the main text indicate good model fits. Overlap between the observed distribution of the response variable (bold-line distributions) and simulations from the posterior predictive distribution (spaghetti lines) indicates the model fits the data well. *Accel born mod* = model testing the relationship between epigenetic age acceleration and birth year, *accel fr mod* = model testing the relationship between epigenetic age acceleration and age at first reproduction, *lrs fr mod* = model testing the relationship between lifetime reproductive success and age at first reproduction.

**Fig. S8.**
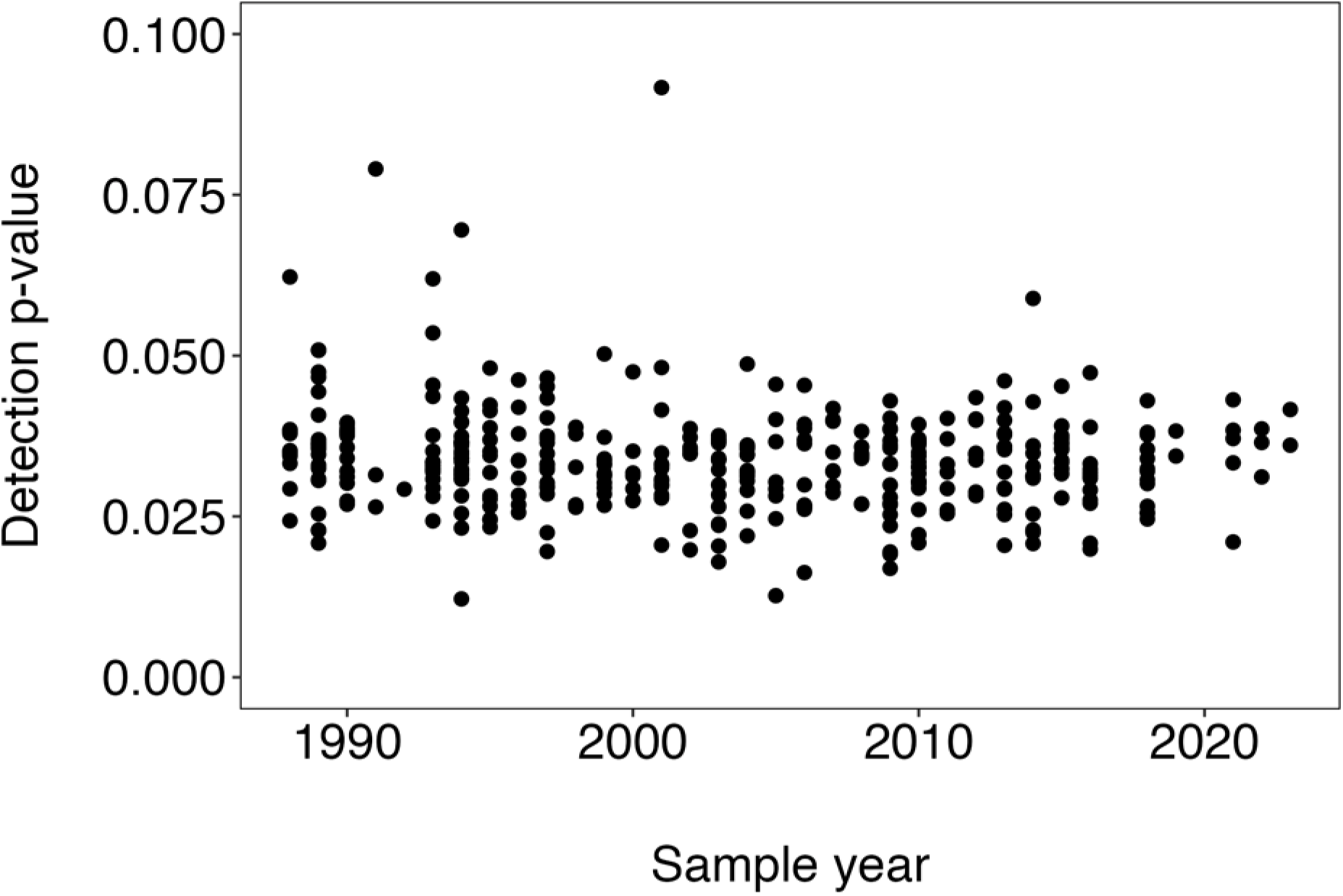
Sample quality is consistent over the range of sample years. Points indicate the detection p-values from samples retained for constructing the epigenetic clock and downstream analysis. Higher detection p-values indicate the CpG sites for the sample showed less discrimination from the background reflectance, indicating DNA methylation signals from the sample are likely unreliable. Plotting the detection p-values over the year of the sample demonstrates quality did not decline with sample age.

**Table S1.**
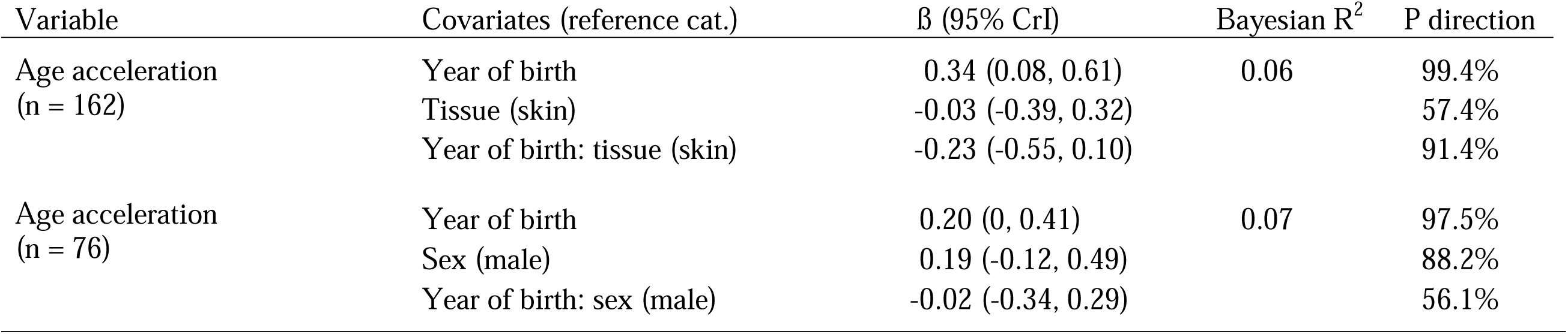
Epigenetic age acceleration over time is consistent between both tissue types and sexes. Coefficients (ß) and 95% credible intervals (CrI) from Bayesian regression models testing relationships between epigenetic age acceleration and year of birth. The two models include terms for tissue type and sex of the sample. The probability of direction (P direction) describes the probability that a coefficient is either positive or negative, expressed as a percentage between 50% and 100%. The Bayesian R^2^ describes the proportion of variance explained by the model. For both models, we used consevative weakly informative priors with mean = 0 and standard deviation = 1. We fit both models using the brms package v2.20.4 in R v4.3.1, with 4 chains and 10,000 iterations including 5,000 warmup iterations.

**Table S2.** There is no genetic variation underlying fitness in the western Hudson Bay polar bear population. Phenotypic mean and estimated random-effect sizes of lifetime reproductive success for western Hudson Bay polar bears using a univariate animal model with a 4,634-individual pedigree. Individuals in the pedigree were documented between 1966 and 2016 in northeastern Hudson Bay near Churchill, Manitoba, Canada. N_ind_ indicates the number of individuals with an estimate of observed lifetime reproductive success, a measure of relative fitness. V_P_ is the total phenotypic variance and is the sum of the variance components. Variance components include V_A_ = additive genetic ‘animal’, V_M_ = maternal ‘dam’ (i.e., the identity of individual’s dam), V_YBirth_ = cohort ‘year of birth’, and V_R_ = residual ‘units’. SD = standard deviation; CI = confidence interval.

| **Trait** | **$N_{\text{Ind}}$** | **$N_{\text{females}}$** | **$N_{\text{males}}$** | **Mean**<br>**LRS**<br>**(SD)** | **$V_P$**<br>**(95% CI)** | **$V_A$**<br>**(95% CI)** | **$V_M$**<br>**(95% CI)** | **$V_{Y\text{Birth}}$**<br>**(95% CI)** | **$V_R$**<br>**(95% CI)** |
| --- | --- | --- | --- | --- | --- | --- | --- | --- | --- |
| Lifetime reproductive success | 456 | 307 | 149 | 4.07<br>(2.57) | 0.16<br>(0.06, 0.29) | 0.008<br>(0, 0.03) | 0.01<br>(0, 0.04) | 0.04<br>(0, 0.07) | 0.11<br>(0.06, 0.15) |

## General

Logistical support for field research was provided by Churchill Northern Studies Centre, Environment and Climate Change Canada, Isdell Family Foundation, Parks Canada, Polar Bears International, Wildlife Media, and World Wildlife Fund. We thank Ian Stirling, Dennis Andriashek and David McGeachy for their invaluable contributions to the western Hudson Bay polar bear program. We also thank C. Kucheravy, E. Karachaliou, E. de Greef, J. Suurväli, and C. Müller of the Population Ecology & Evolutionary Genetics Group at the University of Manitoba, S. Heard, and J.F. Hare, for comments on the manuscript. Finally, we thank A. Shafer, Q. Fletcher, and Bernhard Voelkl and other participants of the 2021 Biomarkers for Stress and Welfare in Wildlife workshop, for sharing code and helpful discussions while preparing the manuscript.

## Author contributions

Conceptualization: LN, ESR, MJJ, CJG

Investigation: LN, HK, OES

Software: LN, LK

Formal analysis: LN, LK

Data curation: LN

Visualization: LN, CS

Resources: ESR, MJJ

Funding acquisition: ESR, MJJ, CJG

Supervision: MJJ, CJG

Writing – original draft: LN, CJG

Writing – review & editing: LN, ESR, BAB, HK, LK, NL, LRR, OES, CS, MJJ, CJG

## Data and materials availability

All data and code are currently available from https://github.com/ljnewediuk/PB_life-history, excluding the pedigree data, which are temporarily embargoed. All data and code will be published as a release in the Zenodo repository upon acceptance.

